# Prolonged impacts of past agriculture and overgrazing on soil fungal communities in restored forests

**DOI:** 10.1101/2020.05.09.086025

**Authors:** Shinichi Tatsumi, Shunsuke Matsuoka, Saori Fujii, Kobayashi Makoto, Takashi Osono, Forest Isbell, Akira S. Mori

**Author notes:** These authors contributed equally to this study (Shinichi Tatsumi, Shunsuke Matsuoka).

## Abstract

Soil fungi can help improve ecosystem restoration, yet our understanding of how they reassemble in degraded land is limited. Here, we studied fungal community structure using DNA metabarcoding in reforested sites following agricultural abandonment and overgrazing. Two treatments, namely ‘reforestation using different numbers of tree species’ and ‘deer exclusion,’ have been applied for multiple decades in the study sites. We found that local fungal richness (alpha diversity) and total fungal richness (gamma diversity) were 1.9–2.9 and 1.3–1.9 times greater, respectively, in reforested stands than in natural forests. These results were regardless of the number of tree species planted in the reforested stands. Conversely, reforested stands had a spatially homogenized community structure with relatively lower degrees of compositional dissimilarity among sites within each stand (beta diversity). These findings were attributable to lower environmental heterogeneity, stronger dispersal limitation, and a comparatively shorter time since the onset of community assembly in reforested stands. Deer exclosures had no detectable effect on fungal community structure. Overall, the agricultural legacy in fungal community structure appears to have persisted for decades, even under proactive restoration of aboveground vegetation. Direct human intervention belowground may therefore be necessary for the recovery of soil biota once altered.

## Introduction

Given the extent of the human-induced damage to the biosphere, ecosystem restoration has become increasingly relevant in today’s world (Hobbs and Harris, 2001). While restoration ecology has typically focused on aboveground vegetation (Brudvig 2011; Perring et al., 2015; Young 2000), less attention has been paid to belowground biota (Kardol and Wardle 2010). The diversity and composition of soil organisms, particularly fungi, can be largely altered in degraded land due to the decreased availability of organic substrates and their symbionts (Kardol and Wardle, 2010). Understanding how soil fungi respond to restoration is of ecosystem-level significance, because they regulate key belowground functions (e.g., litter decomposition and nutrient uptake) and create feedbacks that support the aboveground recovery (Young et al., 2005).

Forest restoration using tree-species mixtures has recently received much social interest (Verheyen et al., 2016; Tatsumi 2020). Plant diversity has often shown to have positive effects on soil fungal diversity in grassland experiments (Milcu et al., 2013; Scherber et al., 2010). In forests, however, experimental studies on the relationships between tree and fungal diversity remain scarce (Weißbecker et al., 2018). Particularly, compared with local fungal richness (alpha diversity), we still know little about the spatial variation in fungal composition (beta diversity), as influenced by tree-species mixing. Assessing both alpha and beta diversity can inform ecosystem recovery not only at local scales but also at larger regional scales that are relevant to forest restoration and policy making.

Aboveground herbivorous can cause significant indirect impacts on soil fungi by altering plant biomass and the production of secondary compounds in foliage and roots (Bardgett and Wardle, 2003; Kardol et al., 2014). Against herbivores, plants can show higher resistance when grown in mixtures than in monocultures, given the associational protection among neighboring plants species (Cook-Patton et al., 2014). The vegetation-mediated effects of herbivores on soil fungi can thus be contingent upon plant diversity. However, no study has yet investigated the possible consequences of such diversity-dependent plant–herbivore interactions on soil fungi by manipulating plant diversity and the presence/absence of herbivores simultaneously.

In this study, we assessed the effects of plant diversity and herbivory on soil fungal communities in a restoration landscape following agricultural abandonment and deer overgrazing. Two treatments were applied in a fully-crossed design: ‘reforestation using different numbers of tree species’ and ‘deer exclusion.’ Our specific objectives were to test (1) whether the community structure of soil fungi, namely their alpha and beta diversity, differ among mixture and monoculture stands as well as nearby natural forests and grasslands, and (ii) whether tree diversity and deer herbivory have individual or interactive effects on soil fungi. Addressing these questions provides a step towards ecosystem restoration grounded on above- and belowground linkages.

## Methods

### Study site

We conducted this study in lowland coastal areas of the Shiretoko National Park, northern Japan (44°08′–11′ N, 145°03′–08′ E, elevation 140–220 m). The area has been designated a World Heritage Site by the United Nations Educational, Scientific and Cultural Organization (UNESCO) on account of it being one of the most species rich, northern temperate ecosystems in the world (http://whc.unesco.org/en/list/1193). The mean monthly temperature ranges from −6.1°C in February to 20.8 °C in August. The mean annual precipitation is 1,149 mm. According to the soil classification system of Japan (Obara et al., 2011) and the Japan soil inventory (https://soil-inventory.dc.affrc.go.jp/), the soil type in our study sites are low-humic allophanic Andosols, which corresponds to Typic Hapludands and Hydric Hapludands in the USDA soil taxonomy (Soil Survey Staff, 2010) (Supplementary Fig. S1).

Approximately 90% of the park’s terrestrial area is covered by pristine natural vegetation, most of which is composed of conifer–broadleaf mixed forests. Parts of the remaining area had been used for agriculture from the early twentieth century until they were abandoned in the late 1960s. Since then, multiple reforestation activities have taken place to restore the ex-arable land to conifer–broadleaf mixed forests (>861 ha; the Shiretoko National Trust Movement Area). Such activities include tree planting with different numbers of tree species and establishment of fences to prevent overgrazing by sika deer (*Cervus nippon yesoensis*). Deer density has increased rapidly from the late 1980s to the late 1990s in the park, with a current density of 6.1–13.6 individuals km^-2^. Currently, the landscape is composed of mosaics of multiple vegetation types, including monoculture and mixture stands, grasslands, and natural forests.

### Study design and sampling

The design and sampling methods are explained in detail in Mori et al. (2016) and Fujii et al. (2017). We used the same study setting and a partly overlapping dataset with Mori et al. (2016) and Fujii et al. (2017). Specifically, we used fungal community data (site × species matrix) and soil environmental data (described in detail below) in this study. In addition to these data, Mori et al. (2016) and Fujii et al. (2017) used data on ecosystem functioning (e.g., litter decomposition).

We used a 4 × 2 factorial design represented by four habitat types and the inside/outside of deer exclosures. The four habitat types were: (i) monoculture stands of *Larix kaempferi*, (ii) mixture stands of *Abies sachalinensis, Picea glehnii*, and *Betula ermanii*, (iii) grasslands dominated by a dwarf bamboo species (*Sasa cernua*) representing a negative control group (i.e., the initial state of restoration), and (iv) natural mixed forests dominated by *A*. *sachalinensis, Quercus crispula*, and *Kalopanax septemlobus* representing a positive control group (i.e., the reference state of restoration).

The deer-exclusion and control sites were established adjacent to each other (i.e., inside and outside of deer fences) in each habitat type. Each site was ~1 ha in size. All the sites were distributed within an area of 2 km × 5 km (Supplementary Fig. S1). Given a restricted number of reforested stands and deer fences in the region, our study consisted of eight sites with one replication for each factorial combination. Reforested stands and deer fences had been established >30 years and ~10 years prior to our field sample collection, respectively (Fujii et al., 2017). The present vegetation inside and outside the deer exclosures show significant structural differences (Fujii et al., 2017; Nishizawa et al., 2016) (Supplementary Fig. S2).

Soil sampling and chemical analyses were conducted as previously described (Fujii et al., 2017; Mori et al., 2016). Briefly, in May 2013, we established three 10 m × 10 m plots in each of the eight factorial combinations. In each plot, we randomly selected three points and collected the topsoil from 0–5 cm depth at each point (i.e., 72 soil samples in total) using a ø 4-cm soil core (DIK-106B; Daiki Rika Kogyo Co., Ltd., Japan). The soil core was washed with ethanol after every use to prevent cross-contaminations. Soil samples were put into separate zip-lock bags, transported from the field on ice, frozen within 3 hours of collection, and kept at −20°C until further analysis.

We measured soil water content, pH, total carbon (C) and nitrogen (N) content, and inorganic N content (ammonium and nitrate). Water content was determined by weighing the soil before and after drying it at 100 °C for 72 h. Soil pH was measured using a pH meter (LAQUAtwin-pH-11B; HORIBA Ltd., Japan) after shaking the soil in water at 1:5 (w/w) for 1 h. Total C and N contents were measured using an organic elemental analyzer (Macro Coder JM1000CN; J-Science Lab Co., Ltd., Japan). Ammonium and nitrate were extracted from soil using a 2-M KCL solution and then measured with an auto-analyzer (AACS-4; BL-TEC Co., Ltd., Japan).

### Molecular analyses and bioinformatics

Molecular analysis and bioinformatics were conducted as previously described (Mori et al., 2016; Fujii et al., 2017) and are explained in detail in Matsuoka et al. (2016b). Briefly, total DNA was extracted from each of the 72 soil samples (0.25 g sample^−1^) using the Soil DNA Isolation kit (Norgen Biotek Corp., Canada). A semi-nested polymerase chain reaction (PCR) was then performed to amplify the nuclear internal transcribed spacer 1 (ITS1) region. The pooled amplicons were sequenced with a GS Junior sequencer (454 Life Sciences, USA).

The reads were clustered with a cut-off sequence similarity of 97% (Osono, 2014) using the Minimus genome assembler (Sommer et al., 2007). Consensus sequences were used as molecular operational taxonomic units (OTUs). A total of 389 OTUs were obtained. For each OTU, taxonomic identification was conducted using the QCauto method implemented in Claident (Tanabe and Toju, 2013). Hereafter, we refer to ‘OTUs’ as ‘species’ for simplicity, bearing in mind that OTUs defined by a fixed sequence similarity do not necessarily represent species in a biological sense. The functional group of each species was determined based on the FUNGuild database (Nguyen et al., 2016) and an intensive literature review (see Tatsumi et al. (2021) for the literature we referred to) <u>(provided as Data S1 for the review process)</u>. See Supplementary Materials for details of molecular analyses and bioinformatics.

### Community structure analyses

We defined fungal alpha diversity as the number of species within a community (i.e., a soil sample) and beta diversity as the extent of community dissimilarity within a treatment combination. Because alpha diversity can potentially be biased by the among-sample variation in sequencing depths (i.e., read counts), we standardized it by using two rarefaction methods, namely sample-size-based and coverage-based rarefactions (Chao and Jost, 2012). We tested the effects of habitat types, deer fences, and their interactions on fungal alpha diversity using the two-way analysis of variance (ANOVA) and Tukey’s HSD test with ‘plot’ as a random effect. The ANOVA and Tukey’s HSD tests were repeated three times for the two rarefied alpha-diversity measures and the non-rarefied alpha diversity (i.e., species richness). In addition to habitat types, deer fences, and their interactions, we included soil properties (pH, total C, total N, C:N ratio, inorganic N, and water content) as explanatory variables in separate ANOVA and linear regression analyses to account for the potential confounding effects these variables could have on alpha diversity. The effects of habitat types, deer fences, and their interactions on soil properties were tested by two-way ANOVA and Tukey’s HSD tests.

To examine the types of rank abundance distribution for each treatment combination, we used four models — preemption, log-normal, Zipf, null models (Wilson, 1991) — which express the abundance of species at rank *r* as functions of *r*. The four models have different functional structures and consist of different numbers of parameters (Wilson, 1991). We estimated the parameters using the maximum likelihood method and calculated the Akaike information criterion (AIC) for each model. Poisson error distribution was used for all models because species abundance was regarded here as a count variable; i.e., the number of sites in which a given species occurred. The model with the lowest AIC was considered the best model for each treatment combination.

The effects of habitat types, deer fences, and their interactions on fungal community composition were tested using the two-way permutational multivariate analysis of variance (PERMANOVA). The compositional dissimilarity between each treatment pair was tested using the pairwise PERMANOVA where *P*-values were adjusted using the Bonferroni correction method. We compared beta diversity among the treatment combinations using the homogeneity of multivariate dispersions test (Anderson, 2006). The communities were ordinated using the nonmetric multidimensional scaling (NMDS). The effects of soil properties on fungal community composition were tested by fitting their vectors onto the NMDS ordination. For all the above community dissimilarity analyses, we used two indices — the Jaccard (Jaccard, 1912) and Raup–Crick indices (Raup and Crick, 1979) — in order to confirm the robustness of the results. The Raup–Crick index is a probabilistic metric which quantifies community dissimilarity that is independent of alpha diversity (Vellend et al., 2007; Tatsumi et al. 2020). The two indices are based on presence/absence data and are thus robust against the possible read-count biases that result from interspecific variation in the ribosomal DNA tandem repeats and from the PCR processes (Toju, 2015). The habitat preferences of each fungal species and functional group were tested based on the association between their occurrence patterns and treatment combinations (De Cáceres et al., 2010). All statistical analyses were implemented in R 3.5.1 (R Core Team, 2018). The R packages used are listed in Supplementary Table S1.

## Results

Fungal alpha diversity was 1.9 to 2.9 times greater in grasslands, monocultures, and mixtures than in natural forests (Fig. 1). We confirmed that the sample-size- and coverage-based rarefactions, as well as the case without rarefaction, yielded qualitatively consistent results; therefore, the results of the last two are provided in Supplementary Fig. S3. The effect of deer exclosures on fungal alpha diversity was not significant (Figs. 1, S3). The interactions between habitat types and deer exclosures also had no detectable effect on alpha diversity (Figs. 1, S3). Soil properties (i.e., pH, total C, total N, C:N ratio, inorganic N, and water content) varied among the treatment combinations (Fig. S4). Habitat type was the most significant predictor of fungal alpha diversity even when we included the soil properties as explanatory variables in ANOVA (Table S2). Regression analyses which controlled for the effect of soil pH (which was found to be a significant variable in ANOVA) also showed that alpha diversity was lower in natural forests than in grasslands (Table S3; Fig. S5).

**Figure 1.**
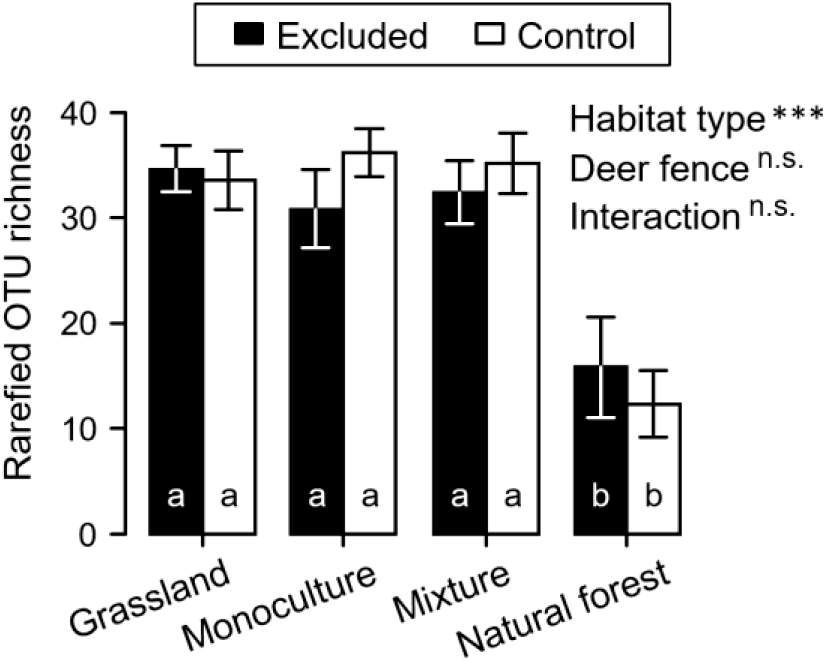
Effects of habitat type and deer fence on rarefied fungal operational taxonomic unit (OTU) richness (i.e., alpha diversity). Results from two-way ANOVA is shown in the upper right; *** *P* <0.001; n.s. *P* ≥ 0.05. Different letters (a and b) in the bars indicate significant differences (*P* <0.05) among treatments (Tukey’s test). Error bars indicate standard errors.

The number of occurrence against species rank decreased less steeply in grasslands, monocultures, and mixtures than in natural forests (Fig. 2). Based on AIC, the preemption model was selected as the best model for the habitats other than natural forests. The Zipf model was selected for natural forests. Gamma diversity (i.e., the total number of species present in each treatment combination) was 1.3 to 1.9 times greater in grasslands, monocultures, and mixtures (>130 species) than in natural forests (≤100 species) (Fig. 2).

**Figure 2.**
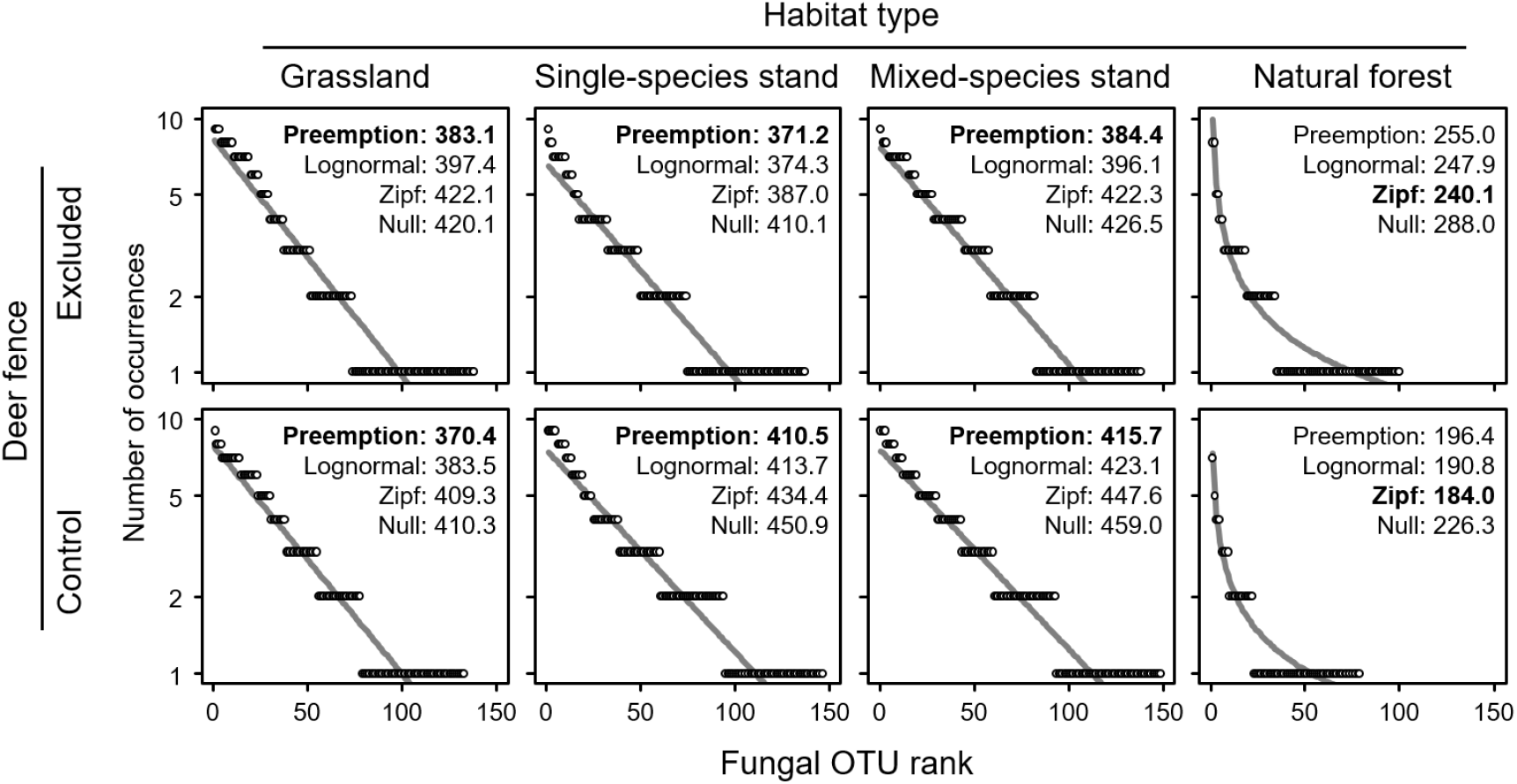
Relationships between the occurrence of fungal operational taxonomic units (OTUs) and their rank. The *y* axes indicate the number of sites in which each OTU occurred within each treatment. The OTUs are arranged in the decreasing order of occurrence frequency. Note that the maximum number of rank (*x* axis) in each panel equals the total number of OTUs observed in each treatment (i.e., gamma diversity). The numbers next to the model names indicate the Akaike information criterion (AIC). Curves show the selected model (shown in boldface), based on AIC, which was fitted to the distribution.

Fungal community composition differed significantly among the habitat types (Fig. 3a, b). We confirmed that the Jaccard and Raup–Crick indices qualitatively yielded the same results; therefore, results from the Raup–Crick index are provided in Supplementary Fig. S6. The effect of deer exclosures on community composition was not significant (Figs. 3a, S6a). The interaction between reforestation types and deer exclosures also had no significant effect on community composition. The soil properties were significantly correlated to community composition (Figs. 3a, S6a). Increases in inorganic N and water content were mainly associated with shifts in fungal composition among habitat types (i.e., arrows roughly paralleled shift direction), whereas soil pH and C:N ratio were associated with community dissimilarity within each habitat type (Figs. 3a, S6a).

**Figure 3.**
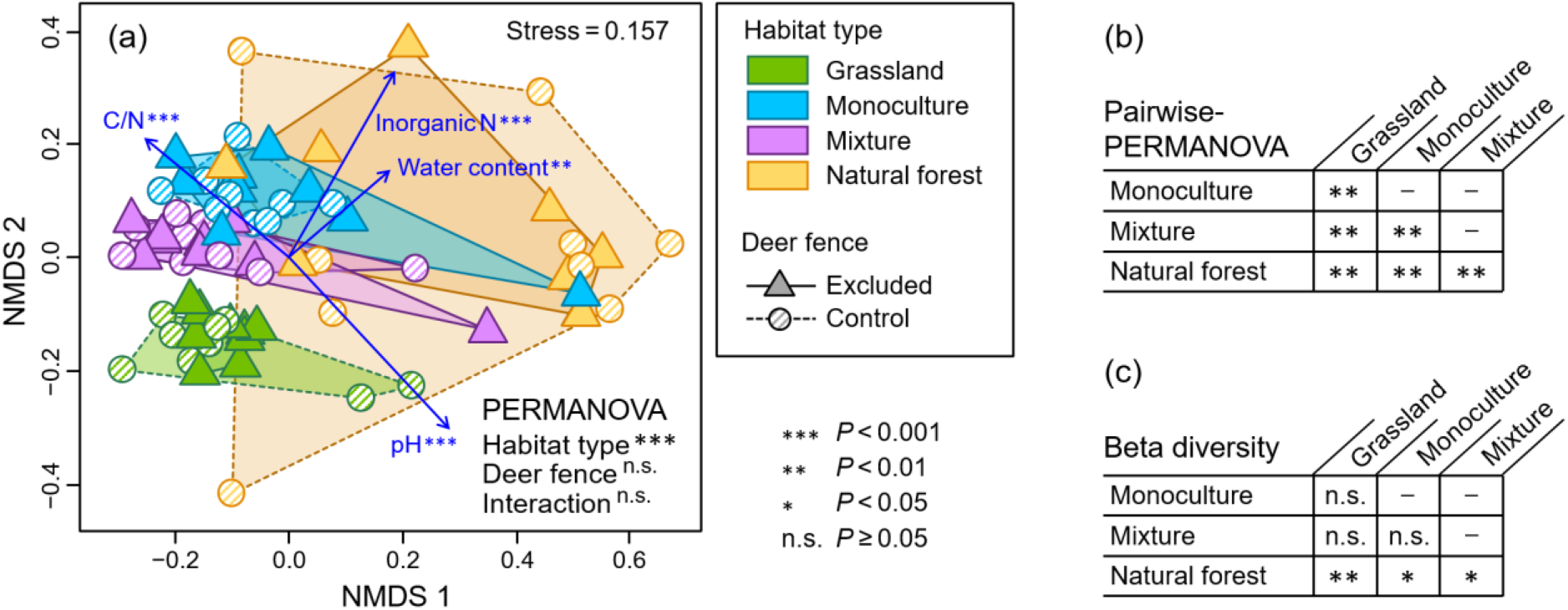
Dissimilarity of fungal communities within and among treatments. (a) Ordination of communities based on nonmetric multidimensional scaling (NMDS) and the effects of treatments (habitat type and deer fence) on community composition tested by two-way permutational multivariate analysis of variance (PERMANOVA). Community dissimilarity was measured using the Jaccard index. Arrows show the associations of soil properties with community composition. (b) Community dissimilarity between pairs of vegetation types. (c) Among-vegetation differences in the size of within-vegetation community dissimilarity (i.e., beta diversity) tested by the permutation test of homogeneity of multivariate dispersion (PERMDISP).

Both Jaccard and Raup–Crick indices showed that the fungal beta diversity (i.e., community dissimilarity within a site) was significantly higher in natural forests compared to the other three habitat types (Figs. 3c, S6c). Note that this pattern should have been statistically independent of the among-habitat variation in alpha diversity (Figs. 1, S3) given the fact that the Raup–Crick index corrects for such variation based on null models (Vellend et al., 2007; Tatsumi et al. 2020).

Most fungal species and functional groups occurred more frequently in certain habitat types (Fig. 4). For example, ectomycorrhizal fungi, ericoid mycorrhizal fungi, and other symbionts that coexist with Ericaceae species occurred more frequently in monocultures and mixtures than in the other two habitat types. Conversely, the majority of fungal species and functional groups occurred at approximately the same frequency both inside and outside deer fences (Fig. 4). Exceptions were *Mortierella* sp. 1 and coprophilous fungi that more often occurred outside than inside fenced exclosures.

**Figure 4.**
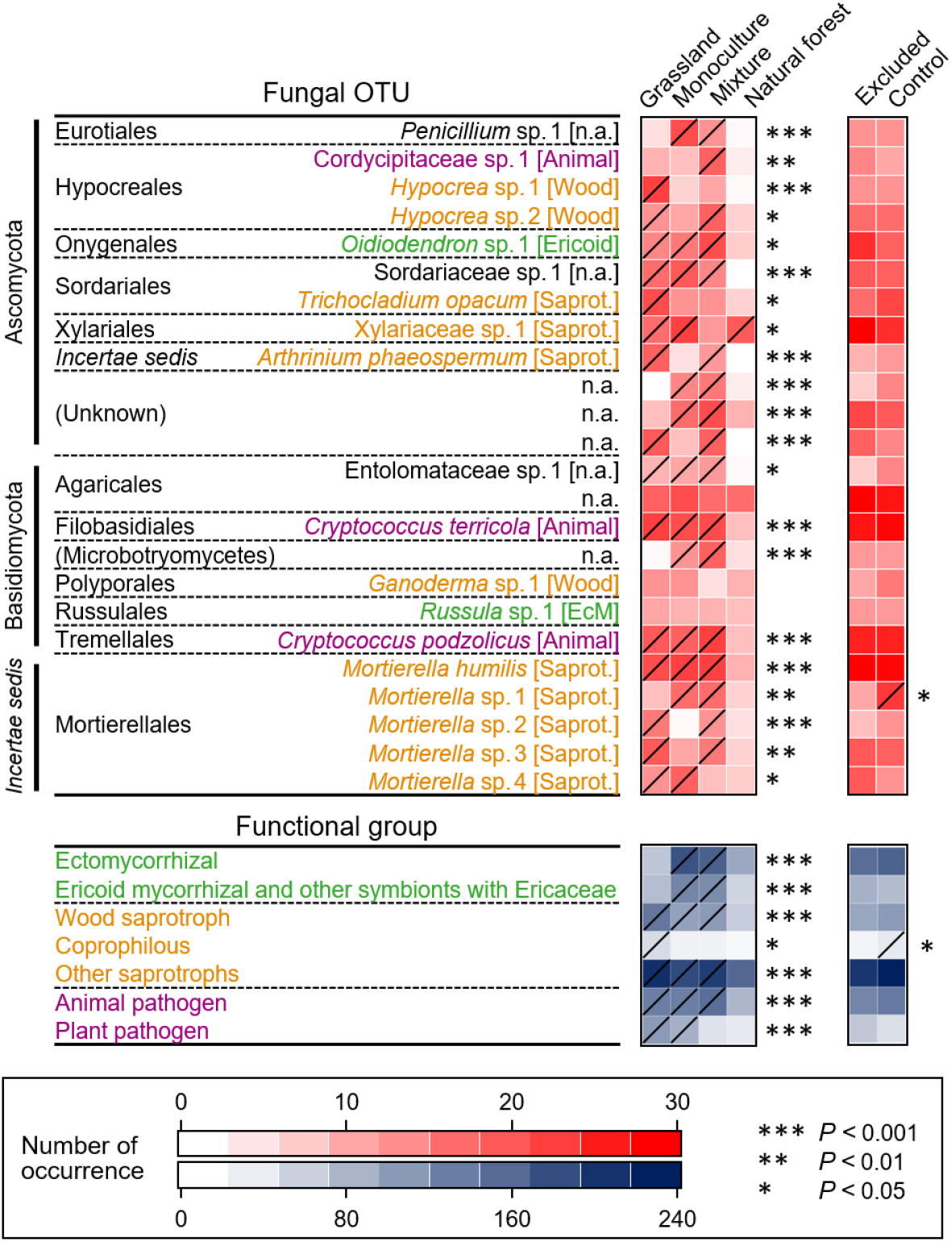
Differences in occurrence frequency of fungal operational taxonomic units (OTUs) and functional groups among restoration treatments. OTUs and functional groups with adequate sample sizes (OUTs occurring at >25 sampling points out of 72 points and functional groups having >3 OTUs) are presented. The transverse lines in boxes indicate treatments in which a given species or a functional group occurred significantly more often than in other treatments.

## Discussion

### Effects of tree diversity and herbivores on soil fungi

Tree planting and herbivore exclusion are among the globally conducted approaches in terrestrial restoration (Kardol and Wardle, 2010; Verheyen et al., 2016). In this study, we tested the effects of these restoration practices on soil fungal communities. Most notably, we found that the local fungal richness (alpha diversity) was 1.9 to 2.9 times greater in grasslands and monoculture and mixture stands than in natural forests (Fig. 1). This result was contrary to the often reported, positive associations between aboveground and belowground biodiversity (Peay et al., 2013; Prober et al., 2015). We also found that the fungal beta diversity and the rank abundance distributions in reforested stands were more similar to those in grasslands than to natural forests, regardless of the number of tree species planted (Figs. 2, 3). The total fungal richness (gamma diversity) was 1.3 to 1.9 time greater in grasslands and reforested stands than in natural forests (Fig. 2). The reforested stands shared more indicator species and functional groups with grasslands than with natural forests (Fig. 4). These results stand in contrast with the fact that aboveground vegetation in our restoration sites is steadily recovering (Nishizawa et al. 2016, Fujii et al. 2017; Fig. S2). Previous studies on naturally regenerating forests showed that historical effects of past land-use activity (e.g., farming and logging) on soil fungal communities can persist for decades (Bachelot et al., 2016; Hartmann et al., 2012). Our results further suggest that, even after decades of proactive aboveground restoration, the soil fungal community structure in reforested stands still differs from that in natural forests.

Deer exclusion had no detectable effect on soil fungal communities at the species level (Figs. 1, 2, 3a). Nevertheless, at the level of functional groups, we found that the coprophilous fungi occurred more frequently outside the fenced exclosures than inside (Fig. 4), a result attributable to the input of deer faeces. Coprophilous fungi were also found most often in grasslands (Fig. 4). This result agrees well with the fact that sika deer prefers grasslands over forests as their habitats (Yabe, 1995). Such deer-induced changes in the functional composition of fungi may, in turn, affect ecosystem nutrient cycling (Kardol et al., 2014). In fact, a faeces decomposition experiment conducted at our study area (Yabe, 1995) found that the faeces decomposed more rapidly (and thus the nutrients released more quickly) in grasslands than forests. This indicates that, in our grassland sites, there is a positive feedback among faeces production, fungal decomposition, and the growth of plants on which deer feed (van der Wal et al., 2004).

Conversely, we found no evidence to support the interactive effects of deer and tree diversity on soil fungi (Figs. 1, 3a). A previous study (Cook-Patton et al., 2014) found a diversity-derived reduction in seedling herbivory by white-tailed deer (*Odocoileus virginianus*) owing to the associational protection of palatable species by unpalatable species. We did not observe such associational effects presumably because sika deer unselectively feed on a wide variety of plants including unpalatable species, especially when its population density is high (Takahashi and Kaji, 2001). Overall, our results indicate that the aboveground plant–herbivore interactions have limited impacts on the species diversity of soil fungi, yet have potential to affect their functional composition and ecosystem functioning.

### Fungal community assembly

Understanding the ecological processes underlying the diversity patterns can provide a critical step towards the development of theory-driven restoration (Mori et al., 2017). Here, based on a conceptual synthesis in community ecology (Vellend, 2010), we discuss three potential assembly processes underlying the lower fungal alpha diversity and the higher beta diversity in natural forests than reforested stands (Figs. 1, 3a, 3c) — that is, ecological selection across space and time, and dispersal limitation.

The first possibility is the selection under different degrees of spatial environmental heterogeneity. Environmental heterogeneity can increase beta diversity (i.e., the dissimilarity among local communities within a given area) through the selection of different species at different sites (Mouquet and Loreau, 2003). The high fungal beta diversity we found in natural forests (Fig. 3a, c) is attributable to the fact that natural forests often have higher habitat heterogeneity compared to grasslands and plantations (Mori, 2011). The greater variation in water content and inorganic N of natural forests compared to grassland and reforested stands (Fig. 3a) also coincides with previous findings that soil properties are often homogenized in ex-arable lands (Bachelot et al., 2016; Fraterrigo et al., 2005). A theory by Kadmon and Allouche (2007) suggests that the environmental heterogeneity can reduce the available habitat size of each species and, thus, could decrease alpha diversity (i.e., local species richness). According to this theory, the low alpha diversity in our natural forests (Fig. 1) could have derived from the high environmental heterogeneity that brought some species to local extinctions.

The second possibility is the time dependency in community response to environmental conditions. Species may not go immediately extinct even when their population growth rates are negative, given the constrained rates of mortality and reproduction. The higher alpha and lower beta diversity in reforested stands compared to natural forests (Figs. 1, 3a, 3c) can be interpreted simply that the processes of local species selection have not yet been completed in reforested stands. Moreover, the long duration of time in natural forests could have allowed the effects of species’ arrival order to be amplified and thereby the communities to diverge (i.e., priority effects; Fukami, 2015). In fact, a multi-year monitoring study of fungal communities (Matsuoka et al. 2016a) found that compositional similarities among communities were largely explained by the closeness in the time of survey. This suggests that the elapsed time after the onset of community assembly can have a major control on fungal community patterns observed in the field. Furthermore, species occurrence patterns in grasslands and reforested stands exhibited the preemption (geometric) distribution (Fig. 2). This type of distribution is often found in the early stages of succession for various taxa, including fungi (Visser, 1995), plants (Whittaker, 1965), and soil invertebrates (Caruso and Migliorini, 2006). On the other hand, the Zipf (power-law) model, which best represented the distribution in natural forests (Fig. 2), indicates that the sites have already been colonized by late-successional specialist species (Gray, 1987). Overall, our results suggest that the fungal species in our reforested sites are still in the course of ecological selection even after decades of tree planting.

The third possibility is dispersal limitation. Despite the small size and immense number of propagules, dispersal limitation is now acknowledged as a crucial determinant of fungal community structure (Peay et al., 2010). Fungal diversity patterns in our study sites (Figs. 1, 3a, 3c) can be explained by dispersal limitation at two conceptual scales: external dispersal from the species pool (as assumed in the mainland–island model) and internal dispersal among local communities (as described in the metacommunity model) (Fukami, 2015). In the former scenario, the low beta diversity of grasslands and reforested stands (Fig. 3a, 3c) can be explained by the limited external dispersal of habitat specialists, which led the communities to become composed of similar suites of generalists, irrespective of local habitat conditions (Vellend et al., 2007). Indeed, a previous study that compared fungal communities in primary forests and artificial pastures (Mueller et al., 2016) found higher dominance of generalist fungi in the pastures. Under the latter metacommunity scenario, maximal alpha and beta diversity is predicted to occur at intermediate and low levels of internal dispersal, respectively (Mouquet and Loreau 2003). This indicates that the low alpha and high beta diversity in our natural forests (Figs. 1, 3a, 3c) resulted from limited within-treatment dispersal. Considering the fact that competition–dispersal trade-offs exist in fungal species (Peay et al., 2007; Smith et al., 2018), it is possible that species with high competitive but low dispersal abilities were only able to subsist in our natural forests.

### Future challenges

We found clear differences in fungal community structure among the treatments. However, uncertainty persists regarding the relationships among some variables we studied. Specially, given the lack of site-level replication for each treatment, we cannot explicitly separate the effects of human activities (i.e., restoration treatments and past agricultural practices) from the among-site environmental variation that might have existed from before (e.g., soil types). For example, we found that the fungal richness and C:N ratio were relatively low in the natural forests (Figs. 1, S3); however, in a strict sense, we cannot tell whether these patterns derive from human activities or the original site environment. In addition, it is possible that some fungal species had been unintentionally co-introduced in the restored stands when the potted tree seedlings were transplanted (Nuñez and Dickie, 2014). Future studies using more strictly-designed experiments (e.g., using multiple site replications, larger sample sizes, and sterilized seedlings) are thus required.

Despite such uncertainties, however, some of our results did indicate that the fungal community structure was in fact driven by the treatments. Most notably, adding the measured soil properties as explanatory variables to the statistical models did not alter our result that fungal richness differed among the habitat types (Table S2). Additional analyses showed that the natural forests had the lowest fungal richness even when we controlled for soil pH which negatively affected the richness (Table S3; Fig. S5). The fact that all the study sites have the same soil type (low-humic allophanic Andosols; Fig. S1) further supports the possibility that the soil environment was homogeneous across the sites prior to the treatments and land-use changes. Moving forward, accumulations of additional field data will allow us to explicitly disentangle the relationships among site conditions, human activities, and fungal diversity.

### Implications for restoration

In this study, we investigated the soil fungal communities in a restoration landscape following agricultural abandonment and overgrazing. We found that aboveground-oriented restoration treatments (i.e., tree planting and herbivore exclusion) do not necessarily translate into the recovery of soil fungal diversity. Our results suggest that in order to enhance the recovery of fungal diversity, direct intervention to the soil, in conjunction with the application of vegetative treatments, may be necessary. For example, supplying organic substrates (e.g., deadwoods) to the soil surface can help create additional habitats for soil fungi (Mäkipää et al., 2017). Especially in restored sites with homogenized community structure like ours (Fig. 3a, 3c), creating mosaics of habitat patches by supplemental substrates may increase the environmental heterogeneity and thus beta diversity. Another commonly applied approach in soil restoration is fungal inoculation, although caution is needed because fungal inoculation can occasionally cause negative impacts on the ecosystem (e.g., due to increased competition among fungal species; Janoušková et al., 2013). In fact, in our study site, natural forests had the lowest fungal alpha diversity (Fig. 1), indicating that simply adding multiple species to degraded land may not necessarily shift the community structure to what is found in natural forests. Rather, it might be effective to selectively inoculate a small number of species that could otherwise not colonize the sites, considering the limited occurrences of habitat specialists in our restored sites (as indicated by low beta diversity; Fig. 3a, 3c). Furthermore, inoculating fungal species in different order at different locations may increase beta diversity via priority effects. We believe that such direct treatments to the soil and belowground biota will allow us to better enhance the recovery of degraded ecosystems.

## Acknowledgements

We thank two anonymous reviewers for invaluable comments, the Shiretoko Nature Foundation for providing logistical support for our field work, and the members of the Shiretoko Biodiversity Evaluation Project for assistance in the field and laboratory work. We are also grateful to Marc W. Cadotte for thoughtful feedbacks, Jia Pu for her support in statistical analysis, Keiichi Okada for his advice on fungal guilds, and Bill Liu for English language editing. ST was supported by the Grant-in-Aids for JSPS Fellows PD (No. 15J10614) and the Young Scientists B (No. 16K18715) from the Japan Society for the Promotion of Science (JSPS). SM was supported by the Grant-in-Aids for Young Scientists B (No. 17K15199) from the JSPS. ASM. was supported by the Mitsui & Co., Ltd. Environment Fund and the Pro Natura Foundation Japan

## Data Archiving Statement

Raw sequence data files are available at the DNA Data Bank of Japan (DRA003024). Fungal community matrix (OTU × sites), species’ functional groups, a list of literature we used to determine the functional groups, and consensus sequences are archived in Figshare: https://doi.org/xxxxxxx.figshare.123456789 (Tatsumi et al., 2021) <u>(provided as Data S1 for the review process)</u>.

