## SupplementaryMaterials for "Prolonged impacts of past agriculture and overgrazing on soil fungal communities in restored forests"

### Supplemental Methods

Total DNA was extracted from each of the 72 soil samples (0.25 g sample$^{-1}$) using the Soil DNA Isolation kit (Norgen Biotek, Thorold, ON, Canada). The polymerase chain reaction (PCR) methods followed those described in Matsuoka et al. (2016). Semi-nested PCR was performed to amplify the nuclear internal transcribed spacer 1 (ITS1) region. In the first PCR, the entire ITS region and 5'-end region of LSU were amplified using the fungi-specific primers ITS1F (5'-CTT GGT CAT TTA GAG GAA GTA A-3') (Gardes and Bruns, 1993) and LR3 (5'-GGT CCG TGT TTC AAG AC-3') (Vilgalys and Hester, 1990). PCR was performed in a 20 µl volume using the KOD FX NEO buffer system (TOYOBO, Osaka, Japan) consisting of 1.6 µl of template DNA, 0.3 µl of KOD FX NEO, 9.0 µl of 2× buffer, 4.0 µl of dNTP, 0.5 µl each of two primers (10 µM), and 4.1 µl of distilled water. The PCR conditions were as follows: an initial step of 5 min at 94 °C, followed by 20 cycles of 30 s at 95 °C, 30 s at 58 °C, and 90 s at 72 °C, and a final extension of 10 min at 72°C. The PCR products were purified using ExoSAP-IT (GE Healthcare, Little Chalfont, Buckinghamshire, U.K.) and diluted by adding 225 µl of sterilized water. The second PCR was conducted using the ITS1F fused with the 454 Adaptor A (5'-CCA TCT CAT CCC TGC GTG TCT CCG ACT CAG-3') and the eight base pair DNA tag (Hamady et al., 2008), and the reverse universal primer ITS2 (5'-GCT GCG TTC TTC ATC GAT GC-3') (White et al., 1990) fused with the 454 Adaptor B (5'-CCT ATC CCC TGT GTG CCT TGG CAG TCT CAG-3'). PCR was performed in a 20 µl volume using the KOD FX Plus NEO buffer system (TOYOBO) consisting of 1.0 µl of template DNA, 0.2 µl of KOD Plus NEO, 2.0 µl of 10× buffer, 2.0 µl of dNTP, 0.8 µl each of the two primer (5 µM), and 13.2 µl of distilled water. The PCR conditions were as follows: an initial step of 5 min at 94 °C, followed by 20 cycles of 30 s at 95 °C, 30 s at 60 °C, and 90 s at 72 °C, and a final extension of 10 min at 72°C. PCR products were checked with agarose gel electrophoresis, purified with ExoSAP-IT, and quantified with Nanodrop. Amplicons were equimolarly pooled into one library and purified using the Agencourt AMPure XP kit (Beckman Coulter, Brea, CA, USA). The pooled products were sequenced following the manufacturer's instructions in a sequencing reaction of a GS Junior sequencer (454

Life Sciences, Branford, CT, USA). Raw sequence data files were deposited in the DNA Data Bank of Japan (DRA003024).

The obtained 101,974 reads were trimmed with a minimum quality value of 27 at the 3' tails (Kunin et al., 2010), and the trimmed reads were sorted into individual samples using the sample-specific tags. The reads with sequence length shorter than 150 bp were excluded, and those longer than 380 bp were shortened to 380 bp by removing bases from the 3'-end. The remaining 67,379 reads were assembled using ASSAMS assembler v0.1.2013.08.10 (Tanabe and Toju, 2013), a highly parallelized extension of the MINIMUS assembly pipeline (Sommer et al., 2007). First, in each sample, reads with 99.5% similarity were assembled and singletons were removed, which potentially represented chimeric sequences and pyrosequencing errors. Reads that were potentially chimeric were discarded using UCHIME 4.2.40 (Edgar et al., 2011). Further, noisy reads were removed using an algorithm in CD-HIT-OTU (Li et al., 2012), after which 39,343 reads remained. The remaining reads were clustered with a cut-off sequence similarity of 97% (Osono, 2014) using the MINIMUS (Sommer et al., 2007). The consensus sequences were used as the molecular operational taxonomic units (OTUs). For each of the obtained OTUs, taxonomic identification was conducted using QCAUTO method implemented in CLAIDENT 0.1.2013.08.10 (Tanabe and Toju, 2013), in which sequences homologous to that of each OTU were searched across the NCBI database. The results from QCAUTO search were then subjected to taxonomic assignment with the lowest common ancestor algorithm (Huson et al., 2007) as implemented in CLAIDENT using the default setting. The OTUs that were not assigned to the kingdom fungi were discarded from further analyses, leaving a total of 389 OTUs and 38,286 reads. The number of reads per plot ranged from 225 to 834 (mean 531.8 ±130.3 SD).

**Table S1.** R packages and functions used for analyses.

| Analysis | Package | Function | Reference |
|---|---|---|---|
| Habitat preference analysis | indicspecies | multipatt() | De Cáceres and Legendre (2009) |
| Sequence rarefaction | iNEXT | estimateD() | Hsieh et al. (2016) |
| Nested ANOVA | nlme | lme() | Pinheiro et al. (2017) |
| Beta diversity test | vegan | betadisper() | Oksanen et al. (2017) |
| Frequency distribution analysis | vegan | radfit() | Oksanen et al. (2017) |
| NMDS implementation | vegan | metaMDS(), envfit() | Oksanen et al. (2017) |

**Table S2.** Effects of treatments and soil properties on rarefied fungal operational taxonomic unit richness as tested by ANOVA.

| Variable | Df | Mean Sq | F value | P value[†] |
|---|---|---|---|---|
| Habitat type (HT) | 3 | 1744.3 | 23.98 | **<0.0001** |
| Deer fence (DF) | 1 | 16.1 | 0.22 | 0.640 |
| HT × DF | 3 | 98.7 | 1.36 | 0.265 |
| pH | 1 | 973.8 | 13.39 | **<0.001** |
| Total C | 1 | 47.2 | 0.65 | 0.424 |
| Total N | 1 | 263.2 | 3.62 | 0.062 |
| C:N ratio | 1 | 22.4 | 0.31 | 0.581 |
| Inorganic nitrogen | 1 | 132.9 | 1.83 | 0.182 |
| Water content | 1 | 70.4 | 0.97 | 0.329 |
| Residual | 60 | 72.7 | | |

[†] Values less than 0.05 are shown in bold letters.

**Table S3.** Effects of habitat types and soil pH on rarefied fungal operational taxonomic unit richness as tested by a linear model.

| Variable | Coefficient | Std. Error | t value | P value † |
|---|---|---|---|---|
| Intercept (grassland) | 184.4 | 41.8 | 4.41 | **<0.0001** |
| Monoculture | −3.8 | 3.0 | −1.26 | 0.211 |
| Mixture | −5.3 | 3.2 | −1.65 | 0.103 |
| Natural forest | −18.5 | 2.9 | −6.33 | **<0.0001** |
| Soil pH | −33.9 | 9.4 | −3.60 | **<0.001** |

† Values less than 0.05 are shown in bold letters.

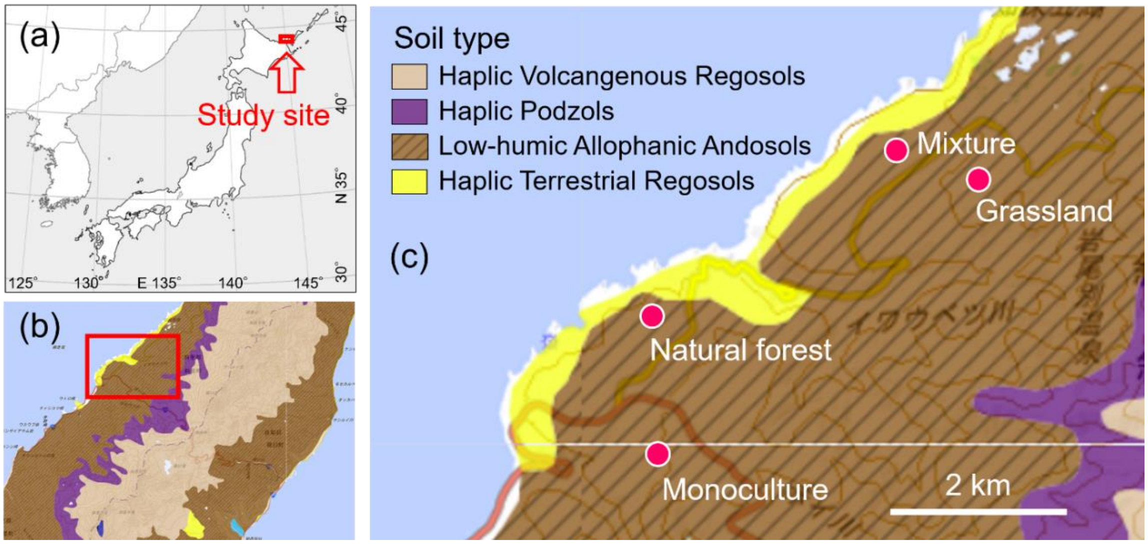


**Figure S1.** Locations and soil types of the study sites. Panels (b) and (c) show the enlarged maps of the areas indicated by red boxes in panels (a) and (b), respectively. The maps in panels (b) and (c) were obtained from the Japan soil inventory (https://soil-inventory.dc.affrc.go.jp/). Soil types followed the comprehensive soil classification system of Japan.

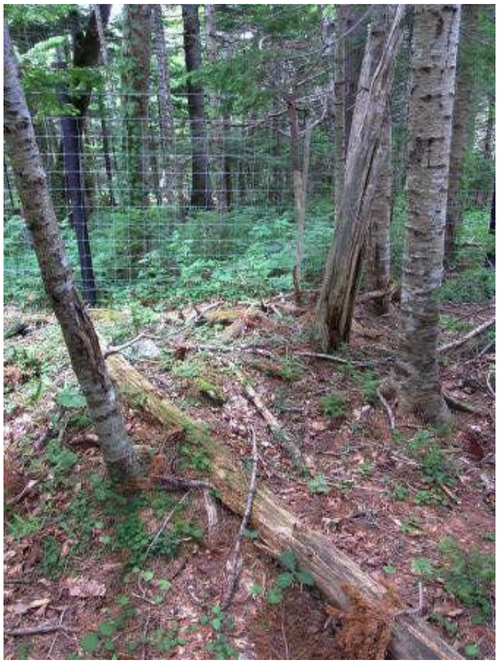

**Figure S2.** A typical example of vegetation inside (at the back) and outside (at the front) of deer exclosures in our study area.

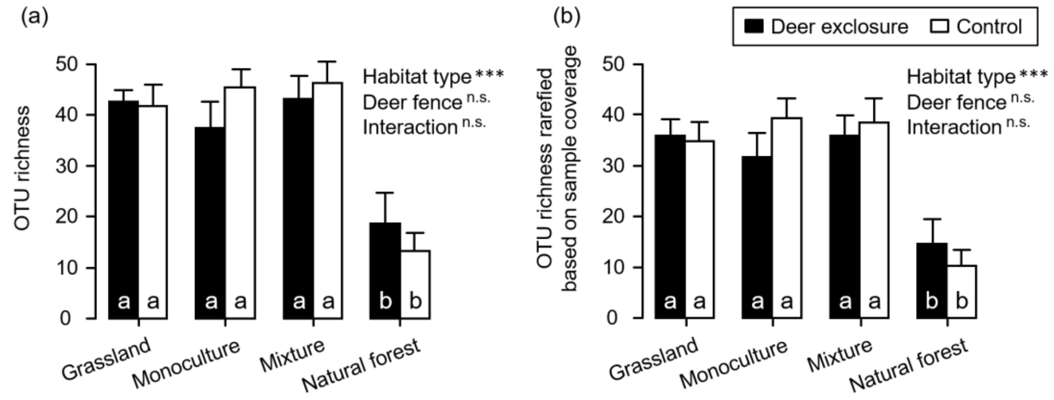


**Figure S3.** Effects of habitat type and deer fence on rarefied fungal operational taxonomic unit (OTU) richness (i.e., alpha diversity). (a) Unrarefied OTU richness. (b) OTU richness rarefied based on sample coverage. Results from linear mixed model analysis is shown in the upper right; *** $P < 0.001$; n.s. $P \geq 0.05$. Different letters (a and b) in the bars indicate significant differences ($P < 0.05$) among treatments (Tukey's test). Error bars indicate standard errors.

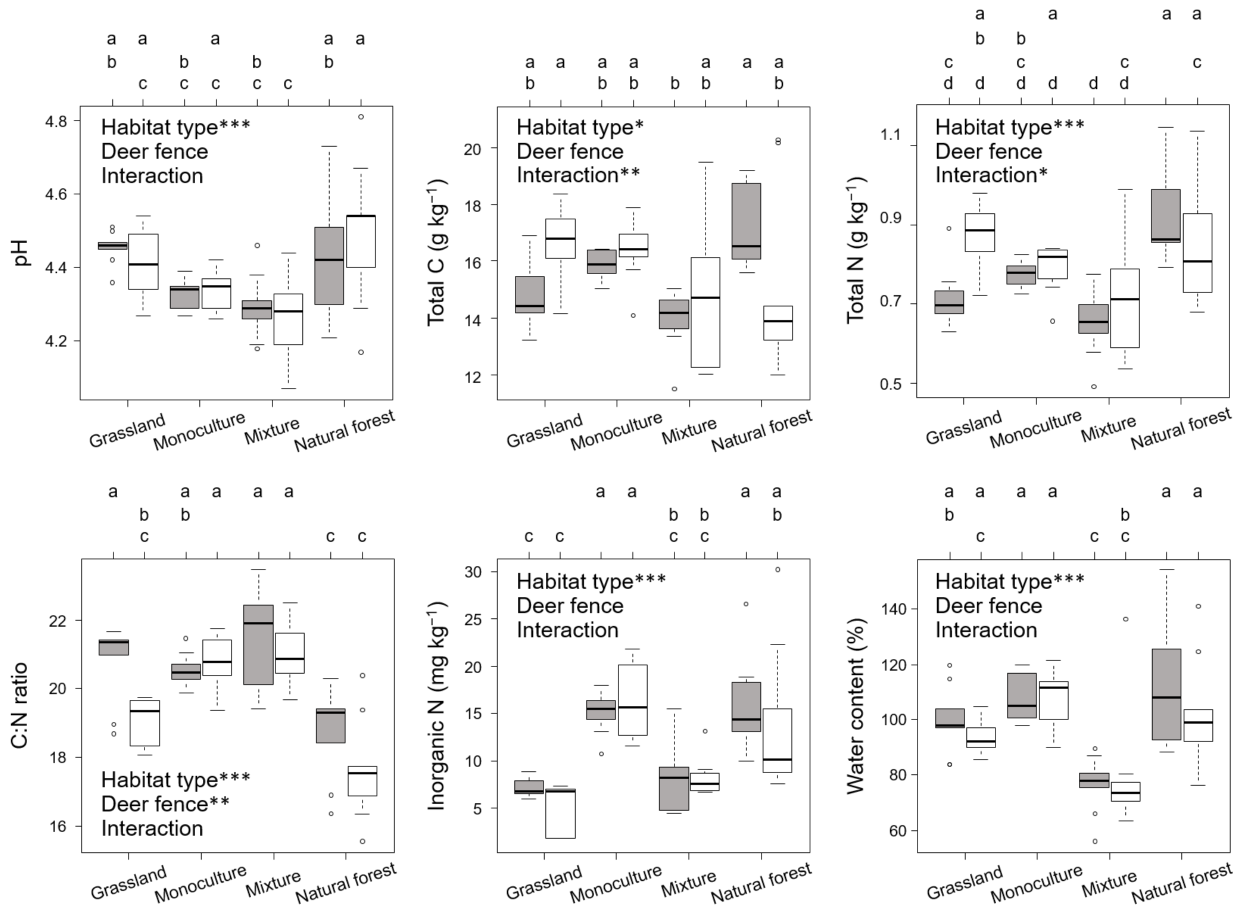


**Figure S4.** Comparisons of soil chemistry among treatments (i.e., habitat types and deer fences). Filled and open boxed show excluded and control sites (i.e., inside and outside deer fences), respectively. Boxes show interquartile ranges (IQR; 25–75th percentile), horizontal lines in the boxes show medians (50th percentile), whiskers show ranges excluding outliers, and circles show outliers which were defined as being outside the IQR by over 1.5×IQR. Asterisks show the results from two-way ANOVA; * $P < 0.05$; ** $P < 0.01$; *** $P < 0.001$. Different letters (a, b, c, and d) at the top of each panel indicate significant differences ($P < 0.05$) among treatment combinations as tested by Tukey's test.

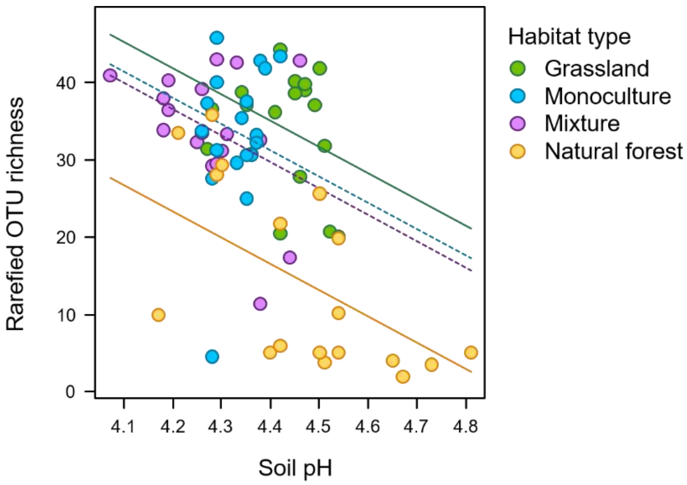


**Figure S5.** Effects of habitat type and soil pH on rarefied fungal operational taxonomic unit (OTU) richness. Lines show the results of regression analyses. Solid lines indicate regression lines that differed significantly to each other (grassland and natural forest). The regression lines that did not differ significantly from the control (grassland) are shown in dashed lines (monoculture and mixture) ($P > 0.05$) (See Table S3).

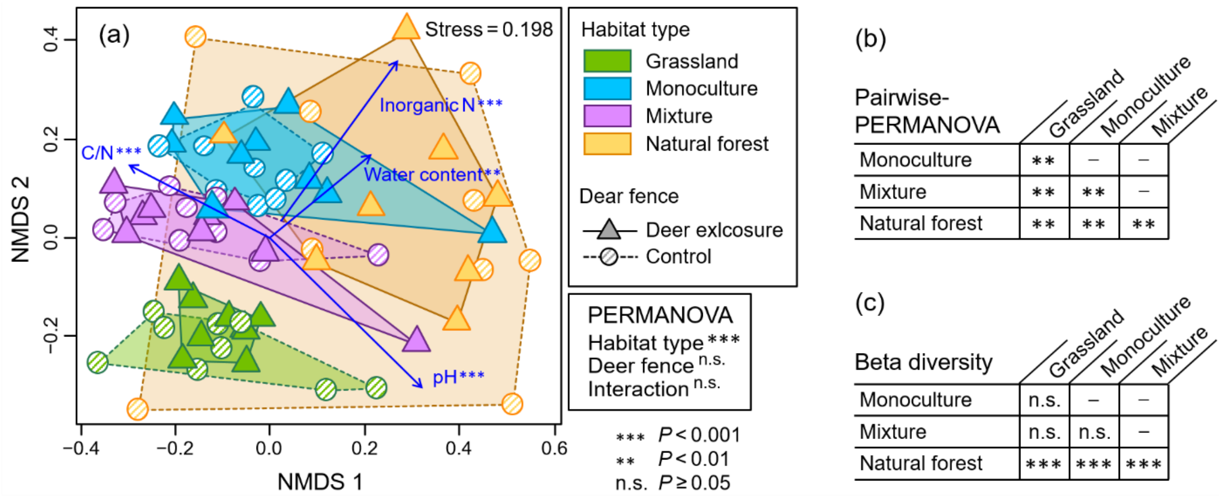


**Figure S6.** Dissimilarity of fungal communities within and among treatments. (a) Ordination of communities based on nonmetric multidimensional scaling (NMDS) and the effects of treatments (habitat type and deer fence) on community composition tested by two-way permutational multivariate analysis of variance (PERMANOVA). Community dissimilarity was measured using the Raup–Crick index. Arrows show the associations of soil properties with community composition. (b) Community dissimilarity between pairs of vegetation types. (c) Among-vegetation differences in the size of within-vegetation community dissimilarity (i.e., beta diversity) tested by the permutation test of homogeneity of multivariate dispersion (PERMDISP).
